# Impact of Essential Genes on the Success of Genome Editing Experiments Generating 3,313 New Genetically Engineered Mouse Lines

**DOI:** 10.1101/2021.10.06.463037

**Authors:** Hillary Elrick, Kevin A. Peterson, Brandon J. Willis, Denise G. Lanza, Elif F. Acar, Edward J. Ryder, Lydia Teboul, Petr Kasparek, Marie-Christine Birling, David J. Adams, Allan Bradley, Robert E. Braun, Steve D. Brown, Adam Caulder, Gemma F. Codner, Francesco J. DeMayo, Mary E. Dickinson, Brendan Doe, Graham Duddy, Marina Gertsenstein, Leslie O. Goodwin, Yann Hérault, Lauri G. Lintott, K. C. Kent Lloyd, Isabel Lorenzo, Matthew Mackenzie, Ann-Marie Mallon, Colin McKerlie, Helen Parkinson, Ramiro Ramirez-Solis, John R. Seavitt, Radislav Sedlacek, William C. Skarnes, Damien Smedley, Sara Wells, Jacqueline K. White, Joshua A. Wood, International Mouse Phenotyping Consortium, Stephen A. Murray, Jason D. Heaney, Lauryl M. J. Nutter

## Abstract

The International Mouse Phenotyping Consortium (IMPC) systematically produces and phenotypes mouse lines with presumptive null mutations to provide insight into gene function. The IMPC now uses the programmable RNA-guided nuclease Cas9 for its increased capacity and flexibility to efficiently generate null alleles in the C57BL/6N strain. In addition to being a valuable novel and accessible research resource, the production of 3,313 knockout mouse lines using comparable protocols provides a rich dataset to analyze experimental and biological variables affecting *in vivo* gene engineering with Cas9. Mouse line production has two critical steps – generation of founders with the desired allele and germline transmission (GLT) of that allele from founders to offspring. A systematic evaluation of the variables impacting success rates identified gene essentiality as the primary factor influencing successful production of null alleles. Collectively, our findings provide best practice recommendations for using Cas9 to generate alleles in mouse essential genes, many of which are orthologs of genes linked to human disease.

## Introduction

The International Mouse Phenotyping Consortium (IMPC) systematically generates and phenotypes mouse lines harboring null mutations in protein-coding genes and prioritizes mouse-human orthologs^1,2^. To produce the majority of its null alleles, the IMPC implemented a deletion strategy to remove a critical region (one or more exons shared by all annotated full-length transcripts that when removed will introduce a frameshift and premature termination codon)^3^. Designs were intended to target protein-coding transcripts for nonsense-mediated decay by introducing a premature termination codon >50-55 nt from the final splice acceptor^4^. While frameshifts can be achieved using single Cas9-mediated double-strand breaks repaired by non-homologous end joining to introduce small insertions or deletions (indels), exon skipping during splicing can restore the reading frame and partial gene function^5–7^. The deletion approach mitigates the risk of restoring frame, simplifies founder screening, genotyping, and quality control (QC), and resembles the embryonic stem cell-based approaches used for decades to produce null alleles, albeit without selection cassette insertion. To evaluate variables affecting mutant mouse production, we analysed data from 4,874 production attempts on 3,973 unique genes from eight different centres (**Supplementary Table 1**) recorded in the IMPC’s production tracking database (GenTaR, formerly iMITS; downloaded 2020 Oct 11, www.gentar.org/tracker/#/). Experimental parameters as well as genomic and biological characteristics of targeted genes were evaluated for their effects on mouse line production success.

## Results

### Experimental parameters affecting mouse line production

Four experimental parameters were assessed for their effects on success: Cas9 delivery method, number of guide RNAs (gRNAs) used, deletion size, and number of founders selected for breeding. We also evaluated the effect of changing parameters in repeated production attempts. Null deletion alleles were generated using Cas9 with guides flanking a critical region containing one or more critical exons^3^. Gene editing reagents were delivered by microinjection (pronuclear or cytoplasmic)^8,9^ or electroporation^10–13^ to target specific genes in C57BL/6N zygotes (**Supplementary Fig. 1**). Among unique gene production attempts (*i.e.,* each gene represented only once; **Supplementary Table 2**), the founder rate, measured as the ratio of founders obtained to the number of embryos treated and transferred, was significantly higher using either cytoplasmic injection or electroporation compared to pronuclear injection of zygotes (**Fig. 1a**; p < 2.22 x 10^−16^, Wilcoxon rank sum test) with no difference between cytoplasmic injection and electroporation (**Fig. 1a**; p=0.26). When we excluded experiments from which no founders were produced, GLT rates by these three delivery methods were all greater than 95% (95.4%, 96.5% and 97.3%, respectively) with no significant difference between them (p > 0.15 in pairwise comparisons with Pearson chi-square).

**Figure 1.**
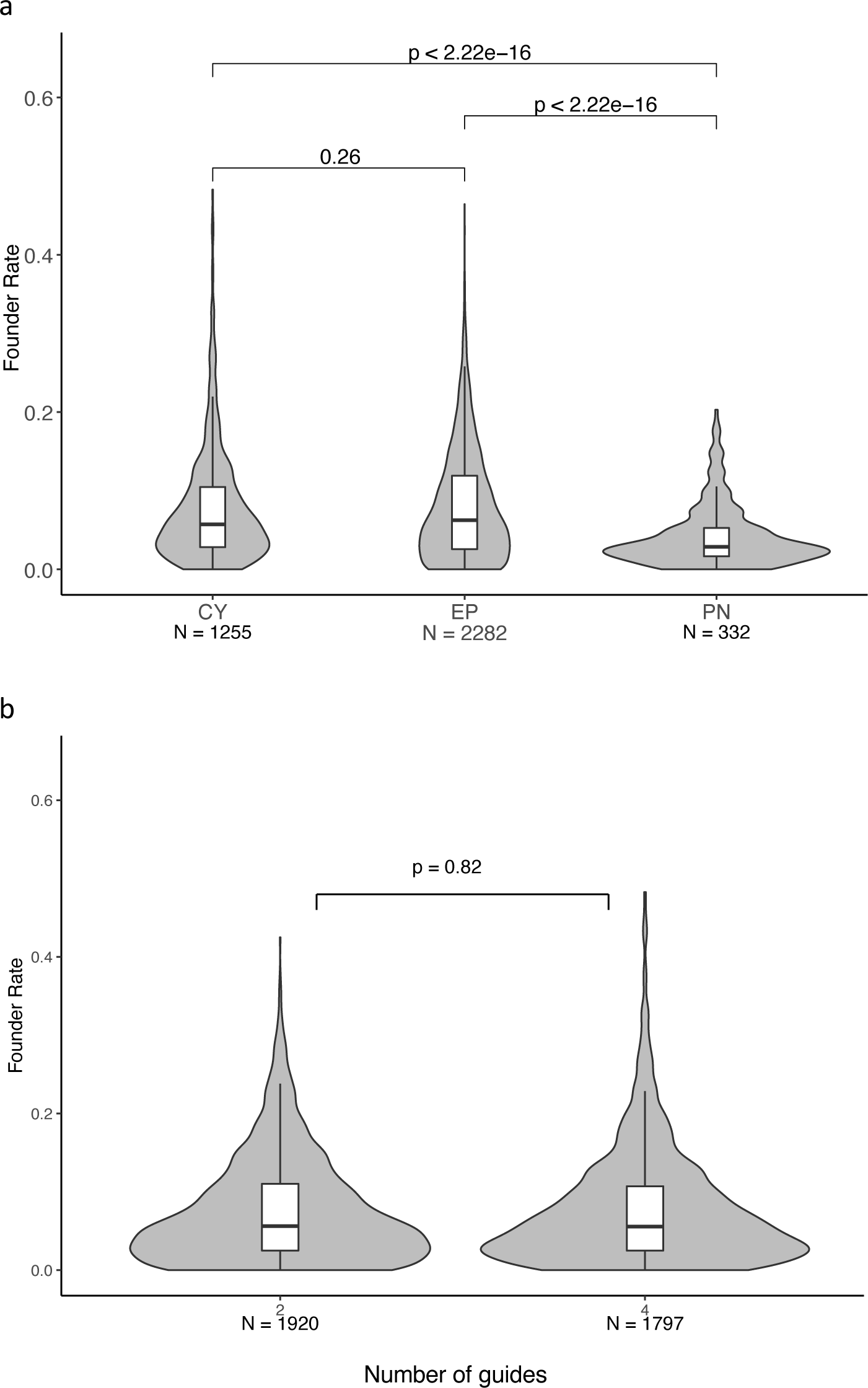
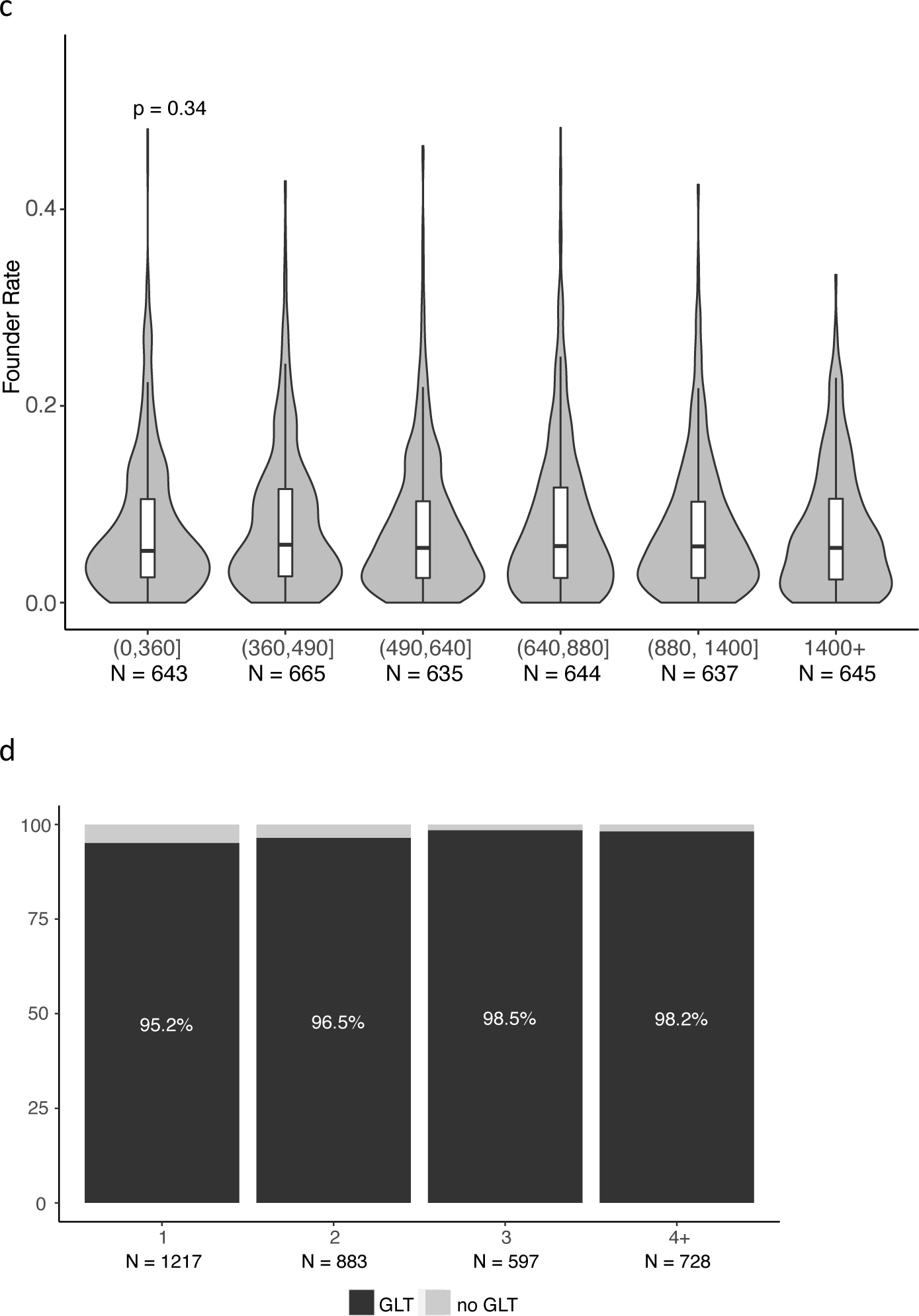
Experimental parameters affecting Cas9-mediated mutant mouse line production. **a**. Founder rates from experiments with different methods of reagent delivery. CY, cytoplasmic injection; EP, electroporation; PN, pronuclear injection. **b**. Founder rates from experiments using two (2) or four (4) guide RNAs designed to produce deletion alleles. **c.** Founder rates from experiments with Cas9 guide RNAs designed to delete different sizes of critical regions (genomic DNA). Each bin has ∼640 unique gene deletion attempts. **d**. Percentage of genes with GLT of the desired deletion allele after breeding one (1), two (2), three (3) or four or more (4+) founders. Pairwise comparison of GLT rates using the Holm method showed a significant difference only when breeding one founder was compared to breeding three or four founders (p = 0.004 for both comparisons). Unique gene attempts are the first attempt with GLT of the desired allele or the last of a set of unsuccessful attempts for each gene. See materials and methods for a complete description of data filtering. GLT, germline transmission.

To mitigate the potential risk of low activity or inactive Cas9-guide combinations, several experiments used four guides, two 5’ and two 3’ flanking the target site. There was no difference in founder rate production (**Fig. 1b**; p=0.82 Wilcoxon rank sum test) or GLT rates of genes edited with two or four guides (GLT 96.8% and 95.7%, respectively, p=0.096 Pearson chi-square). Even with no apparent efficiency gain using additional guides, it is possible that some guides may perform better than others *in vivo* and that inactive or low activity guides may contribute to experimental failure. Using two guides instead of three for four may reduce the small risk of off-target mutagenesis ^14–17^.

To determine if the size of the deleted region influenced success, we partitioned the expected deletion size defined by the maximum distance between flanking guides into six bins with approximately equal numbers of deletion attempts and observed no significant difference in founder rates (p=0.34, Kruskal-Wallis test for comparing medians of six groups; **Fig. 1c**) or GLT rates (p=0.668 Kruskal-Wallis test; data not shown) for deletion sizes below 1,400 bp. The relatively small number of attempts that attempted to delete segments longer than 1,400 bp precludes conclusive statistical analysis of deletions above that size using this dataset, however decreased efficiency with increased deletion size has been reported^18^.

We next assessed whether the number of founders bred affected GLT rate. GLT breeding founders, regardless of production methods used, resulted in efficient germline editing and breeding a single founder provided a >95% chance of success. The overall high GLT rate was marginally improved by breeding up to three founders, but there was no advantage to breeding more than three founders (**Fig. 1d**).

While nearly 70% of first Cas9 production attempts were successful, we wanted to understand why 30% of Cas9 production attempts failed to produce GLT of the desired null allele. Many of these experiments were repeated (with or without changing parameters) and only 26.3% of second attempts and 16.5% of third attempts were successful (**Fig. 2a, Supplementary Table 3**). We analysed the set of repeated attempts for 401 genes to determine what, if any, changes to experimental parameters improved success rates. We compared success rates of consecutive attempts for each gene with repeated attempts to test an association between GLT and each of four experimental parameters (delivery method, number of guides, guide sequence, targeted exon(s)). Changing at least one guide sequence, irrespective of whether the critical region targeted changed, showed a statistically significant improvement in GLT rates (54% with no change to guide sequence *cf*. 68.8% with a changed guide sequence, p=0.01 chi-square; **Fig. 2b**). Changes to other parameters did not significantly affect success rates. A logistic regression model for GLT status conditional on changes in experimental factors also revealed that only changing the guide sequence improved GLT rates (p=0.048, **Table 1**). To evaluate whether these experimental factors interacted to affect GLT rates, we fit an elastic net logistic regression model which performs variable selection and accounts for collinearity through penalization. These regression models provided a 57.58% classification accuracy for GLT status which is only slightly better than a random guess, thus interactions among different parameters could not predict improved experimental success. Therefore, when experiments fail, a significant decrease in success rates after the first attempt should be expected, but changing the guide sequence may improve the likelihood of generating the desired allele compared to making no changes or changing the other parameters examined.

**Figure 2.**
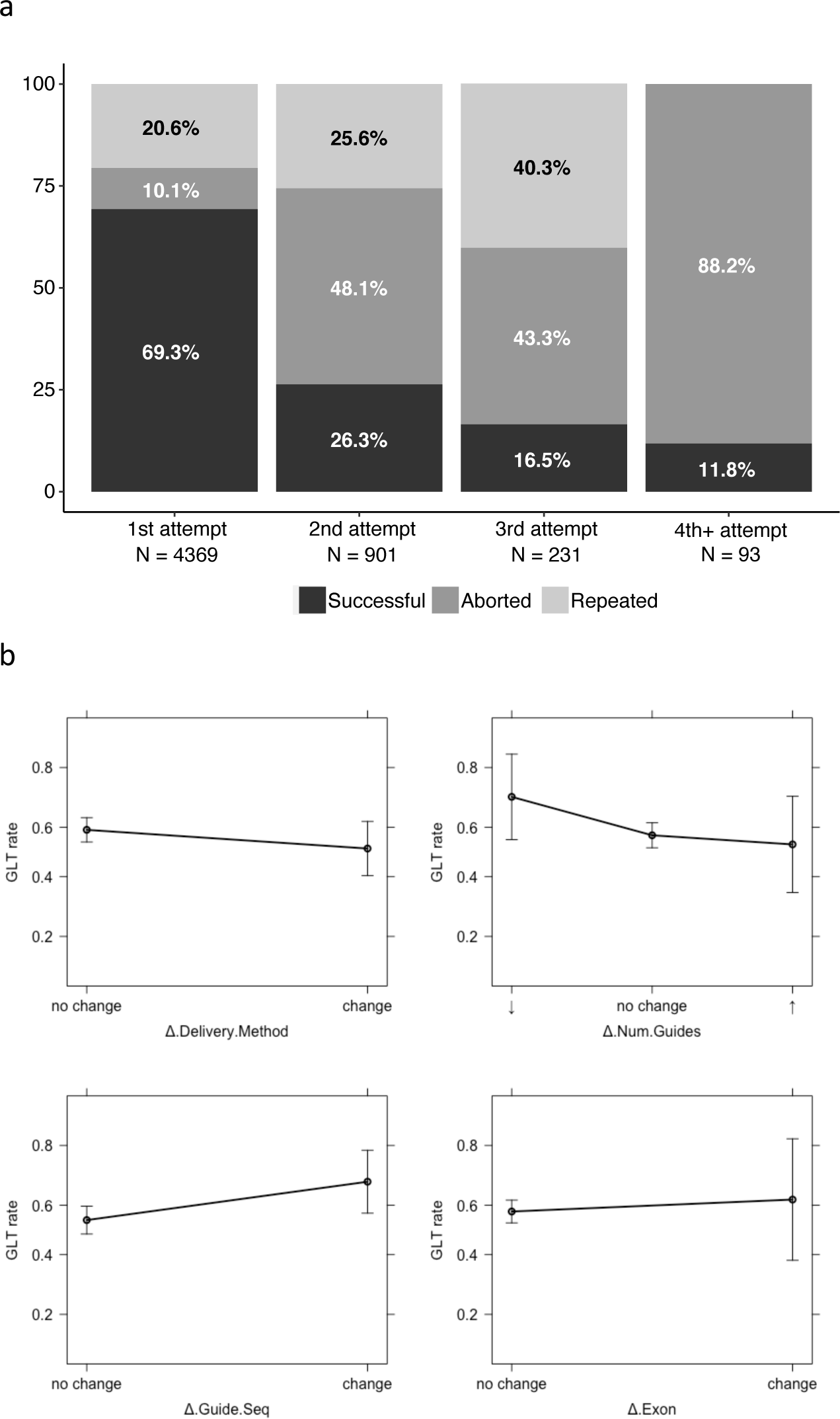
Percentage of genes with GLT of the desired allele. **a**. Percentage of genes with GLT of the desired deletion allele (successful), were abandoned with no additional attempts (aborted), or were repeated in subsequent experiments (repeated). **b**. Effect plots from the linear regression model. A negative slope indicates decreased GLT rates and a positive slope indicates increased GLT rates in subsequent attempts after the indicated parameter changed. See materials and methods for a complete description of data filtering.

**Table 1.**
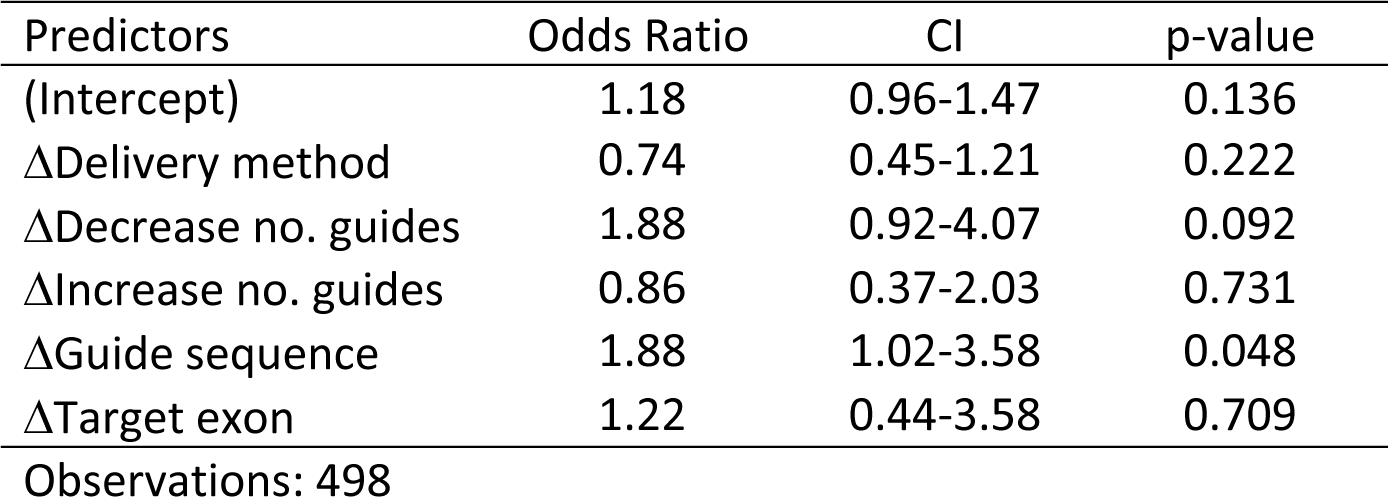
Logistical regression model for GLT status between production attempts conditional on experimental parameters.

| Predictors | Odds Ratio | CI | p-value |
| --- | --- | --- | --- |
| (Intercept) | 1.18 | 0.96-1.47 | 0.136 |
| ΔDelivery method | 0.74 | 0.45-1.21 | 0.222 |
| ΔDecrease no. guides | 1.88 | 0.92-4.07 | 0.092 |
| ΔIncrease no. guides | 0.86 | 0.37-2.03 | 0.731 |
| ΔGuide sequence | 1.88 | 1.02-3.58 | 0.048 |
| ΔTarget exon | 1.22 | 0.44-3.58 | 0.709 |

Observations: 498

### Biological parameters affecting successful mouse line production

In considering biological variables that could influence founder and GLT rates, we hypothesized that targeting essential genes (genes whose function are necessary for cell or animal viability^19^) could negatively affect founder rates due to the high mutagenic efficiency of Cas9. In support of this, the founder rate obtained for cellular non-essential genes was significantly higher than for cellular essential genes (p < 2 x 10^−16^ Wilcoxon rank sum test; **Fig. 3a**). Knockout mouse lines (genes for which knockout alleles were successfully produced) were assessed for homozygous viability of the targeted allele by the IMPC^20^. Alleles identified as homozygous lethal after production, so called lethal genes which include both cellular and developmental essential genes^19^, had significantly lower founder rates than alleles of non-lethal genes (p=2.2 x 10^−11^ Wilcoxon rank sum test; **Fig. 3b**). Similarly, the birth rates for essential and lethal genes during founder production attempts were lower than that for non-essential and non-lethal genes (**Supplementary Fig. 2**). Lower birth rates may reflect a loss of mutant embryos during gestation or shortly after birth due to effects of the introduced mutation. When repeated attempts were classified by gene essentiality^19^ 72.2% of first attempts were successful for cellular non-essential genes compared to 52.6% for cellular essential genes (**Fig. 3c**). In addition, a larger percentage of cellular non-essential genes than essential genes were successful for each subsequent attempt, albeit with decreased success rates for each subsequent attempt for the same gene.

**Figure 3.**
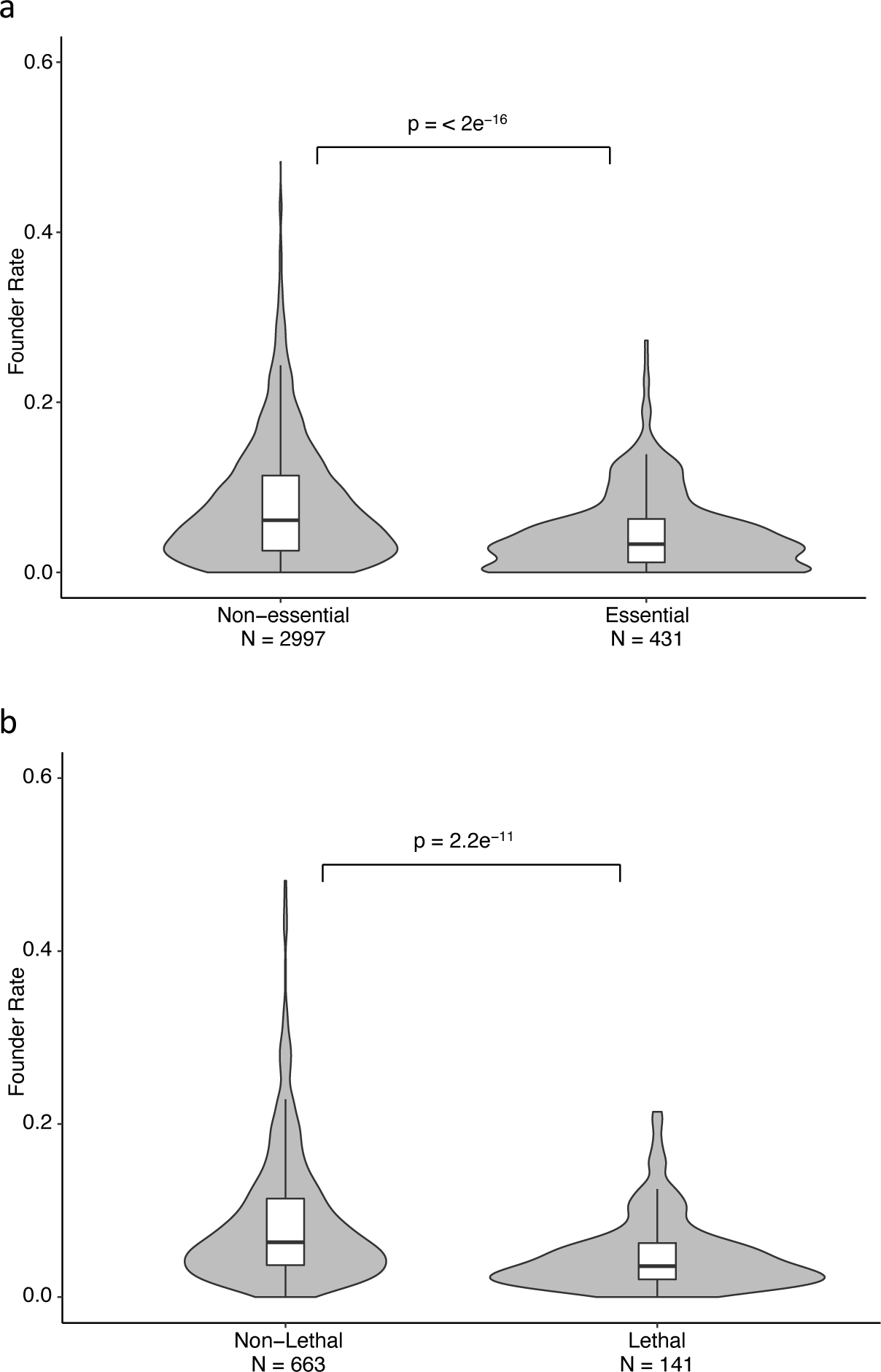
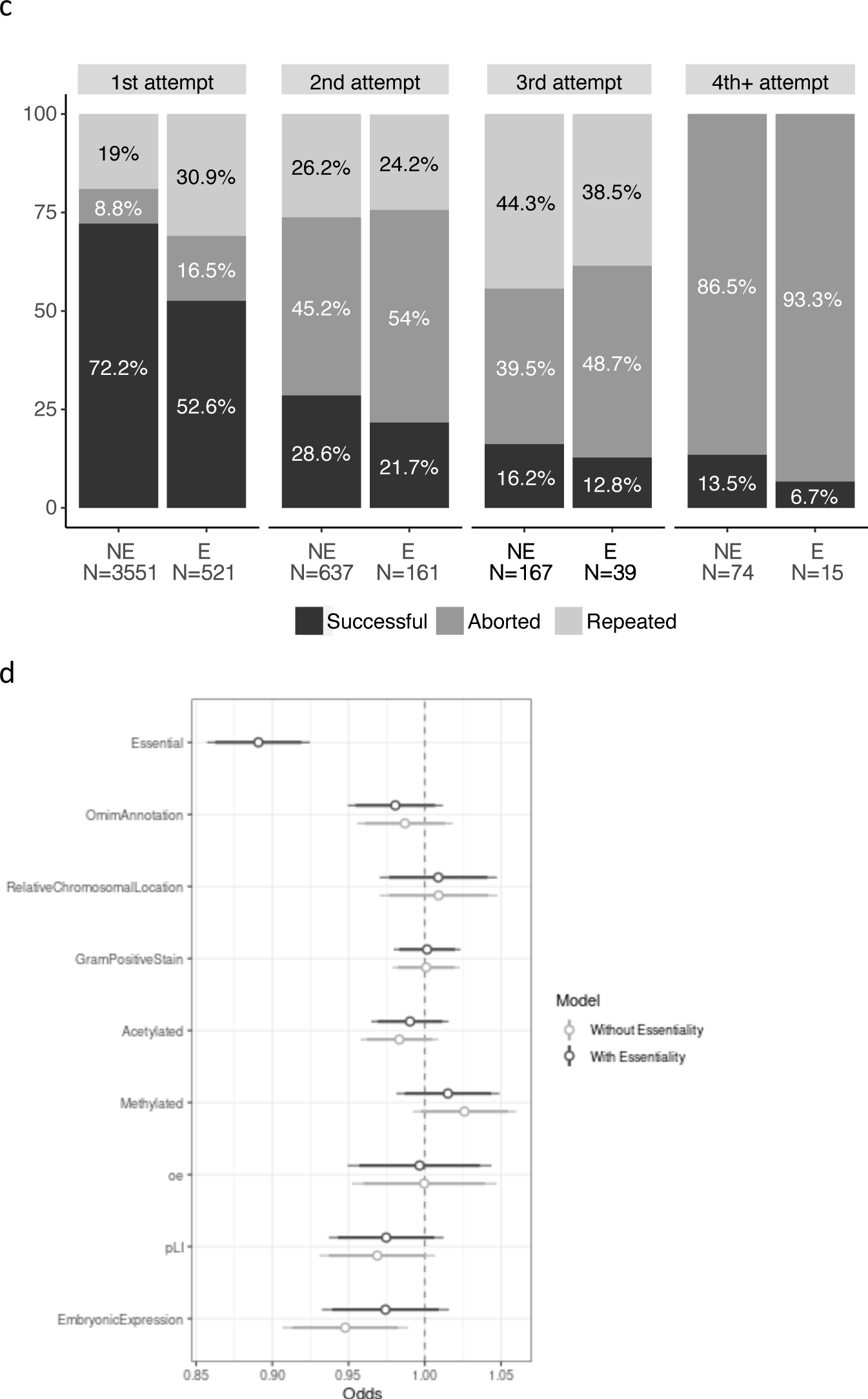
Effects of biological variables on founder and GLT rates for null allele production. **a**. Founder rates of cellular non-essential and essential genes. **b**. Founder rates of homozygous lethal and non-lethal genes. **c**. GLT of null alleles for essential (E) and non-essential (NE) genes with multiple attempts to produce a null allele. **d**. General linear model (GLM) showing the association of each variable with the success of the attempt to generate founders. An odds ratio below 1 is associated with a reduced probability of success, an odds ratio above 1 is associated with an improved probability of success, and an odds ratio of 1 is associated with no effect on success. **Table 2** has the odds ratios and p-values for each variable, with and without essentiality in the model, that assess the significance of the difference of the estimate from zero. **Supplementary Table 5** has the full model output. Each attempt represents a unique gene with the first attempt that successfully generated the desired allele or the last unsuccessful attempt for each gene used for analysis. See materials and methods for a complete description of data filtering.

**Table 2.** General linear model for successful founder production conditional on biological factors annotated for a gene.

| Variable | Without Essentiality |  | With Essentiality |  |
| --- | --- | --- | --- | --- |
|  | Odds Ratio | p-value | Odds Ratio | p-value |
| (Intercept) | 2.50 | $2.0 \times 10^{-16}$ | 2.52 | $2.22 \times 10^{-16}$ |
| Essential | NA | NA | 0.89 | $2.06 \times 10^{-11}$ |
| Embryonic expression | 0.95 | 0.011 | 0.97 | 0.219 |
| pLI score | 0.97 | 0.101 | 0.97 | 0.183 |
| o/e score | 1.00 | 0.987 | 1.00 | 0.887 |
| Chromosome position | 1.01 | 0.647 | 1.01 | 0.653 |
| Acetylated gene | 0.98 | 0.193 | 0.99 | 0.450 |
| Methylated gene | 1.03 | 0.137 | 1.01 | 0.383 |
| Gram positive stain | 1.00 | 0.938 | 1.00 | 0.891 |
| OMIM annotation | 0.99 | 0.412 | 0.98 | 0.220 |

Observations: 3209

There was a small, but not statistically significant, difference in GLT rates between cellular non-essential and cellular essential genes (96.8% and 95.2%, respectively; p=0.15 Pearson Chi-square) among experiments that generated founders. This may be a result of fewer founders obtained and therefore fewer bred for cellular essential genes compared to non-essential genes. Alternatively, this may have occurred if haplo-insufficient genes were targeted and produced mosaic founders that could not propagate mutant progeny that reached genotyping age.

To assess the influence of additional biological variables on founder and GLT rates, we applied a general linear model (GLM) to test the association of several factors including embryonic expression (GEO GSE11224)^21^, observed/expected loss-of-function (o/e) score, probability of loss-of-function (pLI) score^22^, chromosome position, histone methylation and acetylation (as a proxy for chromatin accessibility), cytogenic banding (a second proxy for chromatin accessibility), and human ortholog disease annotation in the Online Mendelian Inheritance in Man (OMIM) database ^23^ (**Supplementary Table 4**). When gene essentiality was excluded from analysis, only embryonic expression was significantly associated with low founder rates (odds ratio (OR)=0.95, p=0.011). However, when essentiality was included, it was the only predictor of experimental failure (OR=0.89, p=2.06 x 10^−11^; **Fig. 3d**, **Table 2, Supplementary Table 5)**. The proportion of essential genes in each experimental parameter grouping did not vary in a way that may have confounded the results of those analyses (**Supplementary Tables 6 and 7**).

After successful production of a founder, subsequent GLT rates were very high (**Fig. 1d**). Among 74 genes for which the reason for GLT failure was readily determined the majority failed GLT because founders died before breeding, founders failed to produce progeny, or founders propagated only wild-type progeny that reached genotyping age (**Fig. 4**). The remaining genes failed primarily because the desired null allele could not be validated in N1 progeny (*e.g.,* partial deletion or no frameshift recovered). Infertile founders and transmission of only wild-type progeny might be due to founder mosaicism with only wild-type cells contributing to the germline or mutation effects on gametogenesis with only wild-type cells able to produce functional germ cells. Moreover, transmission of only wild-type alleles could indicate the presence of haplo-insufficient alleles for some genes causing mutant progeny to die before genotyping. These data support the hypothesis that a substantial subset of mutations may fail GLT due to negative effects on viability or fertility and are consistent with the significant effect of gene essentiality on founder production (**Table 2**).

**Figure 4.**
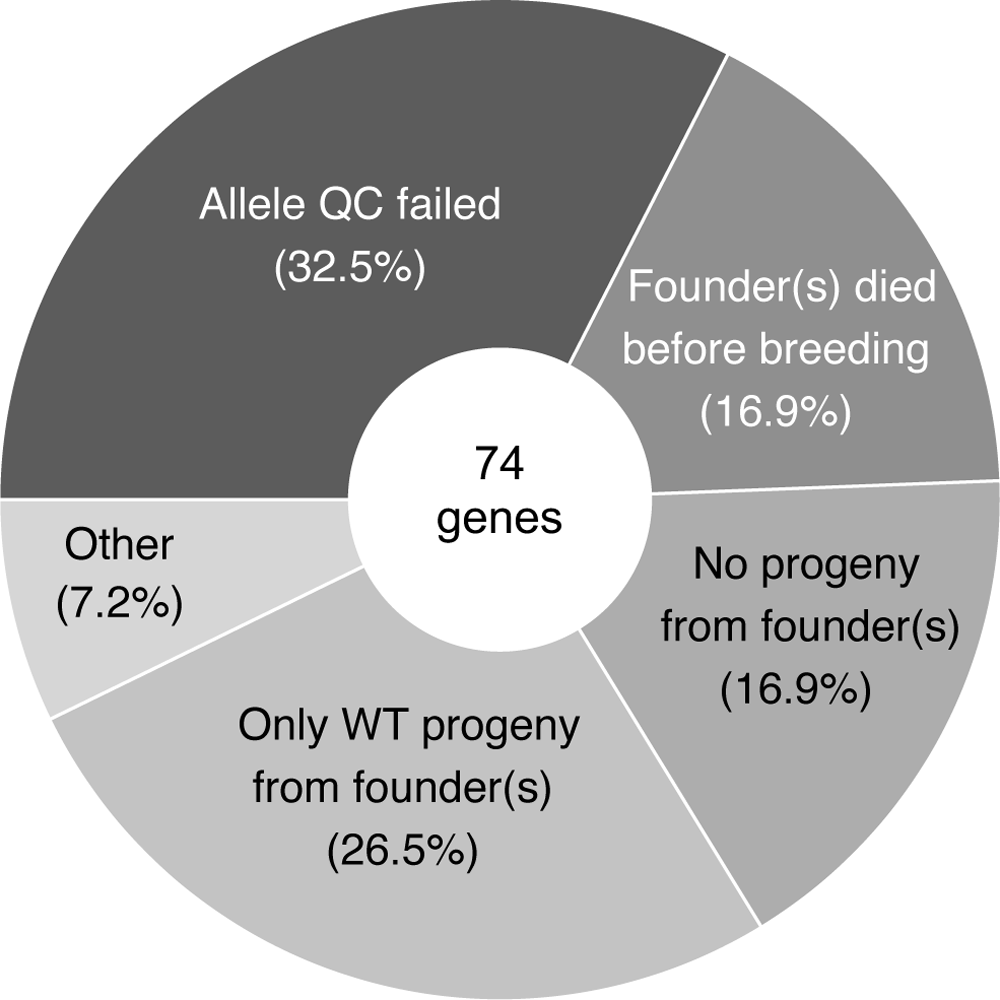
Summary of reasons founders failed to transmit a quality-controlled deletion allele to subsequent generations to establish a knock-out mouse line.

## Discussion

The analysis of our large, multi-centre dataset identified several variables that affect Cas9 mutagenesis success and provides the basis for recommendations for genome editing in mouse zygotes. For deletion alleles, Cas9 ribonucleoprotein (RNP) electroporation is the most accessible and scalable method, providing equivalent performance to cytoplasmic microinjection of Cas9 mRNA and gRNAs. GLT rates are high for Cas9 generated founders and there is no apparent advantage to breeding more than three founders. Most attempts are successful the first time, but for attempts that fail to produce founders, changing guide sequences may be beneficial when repeating production for a given gene. The activity of Cas9-guide RNA combinations can be assessed *in vivo* (*e.g.*, by zygote treatment, *in vitro* culture, and blastocyst genotyping^24^). Exploring the known biology of the gene, such as predicted effects on phenotype (*e.g*., disease association of human ortholog), viability, and/or fertility as well as essentiality scores from publicly available sources (as in ^19^), can assist with experimental design. Successful targeting of essential genes may require methods that promote founder heterozygosity (*e.g.,* lower Cas9 concentrations) or mosaicism (*e.g.,* delivering reagents to one blastomere of a 2-cell embryo by microinjection). In some cases, conditional alleles^25,26^ may be required instead of constitutive knockout alleles to generate a mouse line in which homozygous loss of function may be examined in specific tissues or developmental stages.

The effect of cell essentiality or lethality can apply to generating lines with disease-associated variants such as single nucleotide variants. Given the high proportion of essential genes among human disease genes^19,27^, generating mouse models of human disease may require preserving one copy of the wild-type allele, that is producing heterozygous founders or conditional variant alleles, to establish a new mouse line. Overall, the use of Cas9 is a robust and flexible method to generate gene-edited mouse lines. Improving model production success rates by applying our recommendations can reduce the number of animals used to generate new models, consistent with the ethical principles of the 3R’s^28^.

## Materials & Methods

### Mouse strains

All null allele mouse lines were produced in the C57BL/6N strain background available from Charles River or the Jackson Laboratory (**Supplementary Methods Table 1**). All live animal protocols conformed to the applicable standards for the ethical use of animals in research at the respective facilities with detailed ethics statements found in **Supplementary Methods Table 1** for each production centre. Animal welfare was regularly monitored.

### Allele design

Null alleles were designed such that the mutations resulting from Cas9 endonuclease activity and double-strand break (DSB) repair caused a reading frameshift in protein-coding transcripts and introduced a premature stop codon. This required the identification of a critical region for each targeted gene. A critical region was defined as one or more exons that when frameshifted or deleted resulted in a frameshift in the open reading frames of all known full-length protein-coding transcripts (per Ensembl build 38)^3^. For the majority of designs, the premature stop codon was predicted to be in the first half of the protein-coding open reading frame and to target transcripts for nonsense-mediated decay^4,29^. Such alleles are considered presumptive nulls. Three major categories of alleles were evaluated in this study. Exon deletion (exdel) alleles resulting from NHEJ-mediated repair of Cas9-induced DSBs flanking the exon(s) within a gene’s critical region. Intra-exon deletion (intra-exdel) alleles resulting from repair of Cas9-induced paired DSBs within a single exon in the target gene’s critical region, used for example, when all exons of a gene are in the same frame precluding an exon deletion strategy or for single exon genes. Inter-exon deletion (inter-exdel) alleles resulting from repair of Cas9-induced paired DSBs in two or more different (often sequential) exons in a gene’s critical region, used for example, to delete a functional domain that spans multiple exons or when specific gRNAs in appropriate intron locations could not be identified. Alleles that did not result in a frameshift failed quality control (QC) metrics and were not maintained. Exceptions were made when the deletion was still in frame, but either removed critical protein domains (*e.g*., zinc fingers from a zinc-finger protein) or a substantial fraction of the protein-coding sequence; these alleles were also deemed to be presumptive null allele.

### sgRNA selection

Guide RNA (gRNA) spacer sequences were selected using either the CRISPR design tool^30^, the Wellcome Sanger Genome Editing (WGE) Tool^31^, CRISPOR^32^, CRISPRTools^33^, CHOPCHOP^34^, or FORCAST^35^. Suitable gRNA spacer sequences were selected to minimize predicted off-target mutagenesis using specificity scores >65 when available and/or sequences with at least 3 mismatches for all predicted off-target sites. Multiple guides were used to generate deletion alleles, with either two, three, or four guides (2G, 3G, or 4G, respectively) with two or more guides flanking the target critical region. In the 3G approach, the middle guide could be within an exon and result in the deletion in conjunction with either the upstream or downstream guide, removing either the splice acceptor or donor from the critical region, resulting in mis-splicing and the introduction of a frameshift.

### sgRNA synthesis

sgRNAs were synthesized by either subcloning sgRNA spacer sequences and *in vitro* transcription (plasmid-IVT), PCR and *in vitro* transcription (PCR-IVT)^36^, gBlock synthesis and *in vitro* transcription (gBlock-IVT), or by primer extension and *in vitro* transcription (PE-IVT)^36^. Alternatively, sgRNAs were purchased from commercial suppliers. See **Supplementary Methods Table 1** for centre-specific reagent details.

#### PCR-IVT

DNA templates for PCR-IVT were produced using overlapping oligonucleotides in a high-fidelity PCR reaction^37^ or using a plasmid template (Addgene #42230^38^) and appropriate primers^36^. PCR amplicons were purified using the Monarch PCR & DNA cleanup kit (New England BioLabs T1030) or the QIAQuick PCR purification kit (Qiagen 28104) and used as a template for *in vitro* transcription of the sgRNA with the T7 MEGAshortscript™ Kit (ThermoFisher AM1354).

#### Plasmid-IVT

Overlapping oligonucleotides with *Bsa*I appendages to facilitate standard sticky ended cloning into a T7 expression plasmid (a kind gift from Sebastian Gerety, based upon^39^) were purchased annealed. Alternatively, annealed oligonucleotides were cloned into plasmid DR274 (Addgene #42250^40^). Plasmid DNA was extracted using the QIAGEN Plasmid Plus 96 Kit (Qiagen 16181) and guide cloning confirmed by Sanger sequencing. The DNA was then linearized and used as a template for T7 RNA *in vitro* transcription using the T7 MEGAshortscript™ Kit (ThermoFisher AM1354) or Thermo T7 RNA polymerase (TOYOBO, TRL-201).

#### gBlock-IVT

sgRNAs were synthesized directly from gBlock® DNA (Integrated DNA Technologies) templates containing the T7 promoter using the HiScribe™ T7 High Yield RNA Synthesis Kit (New England BioLabs E2050) following manufacturer’s instructions for sgRNA synthesis.

#### PE-IVT

The EnGen sgRNA Synthesis Kit (New England BioLabs E3322) was used for PE-IVT per the kit protocol, but with incubation at 37°C for 60-90 minutes prior to DNAse treatment. For some 3G and 4G designs, up to two primers (*e.g.* both upstream gRNAs or both downstream gRNAs) were pooled at appropriate final concentrations before PCR or PE-IVT. After *in vitro* transcription, sgRNA was purified using the RNA Clean & Concentrator-25 (Zymo Research R1017) or the MEGAclear Transcription Clean-Up Kit (ThermoFisher AM1908). All samples were analyzed by Nanodrop to determine A260/280 and A260/230 ratios (≥1.9 to pass quality control). The integrity and size of sgRNA was assessed by agarose gel electrophoresis, Agilent Bioanalyzer, Agilent RNA Tape or the Qiaxcel Advanced System (RNA QC V2.0).

Synthesized sgRNAs were stored at −80°C in elution buffer or stored as ammonium acetate precipitates in ethanol at −20°C. Before use, sgRNAs were either thawed on ice or pelleted, air dried, and resuspended in RNAse-free MI buffer.

### Cas9

Cas9 mRNA was purchased (**Supplementary Methods Table 1**) or transcribed in-house^41^. Cas9 protein was purchased from commercial suppliers. See **Supplementary Methods Table 1** for centre-specific reagent details.

### Injection mix preparation

Injection mixes were prepared essentially as previously reported^36^ with or without filtration prior to injection. For mRNA microinjection, injection mixes consisted of Cas9 mRNA and sgRNA in microinjection buffer (**Supplementary Methods Table 1)**.

Concentrations for each production attempt are shown in **Supplementary Table S8**. For Cas9 protein microinjection, Cas9 ribnucleoprotein (RNP) complexes were produced by mixing the Cas9 protein with sgRNA at 5X the concentration shown in **Supplementary Methods Table S1** in RNP injection buffer and incubating at 37°C or room temperature for 10 minutes. The RNP mix was then diluted with 4 volumes of RNP injection buffer prior to injection. See **Supplementary Methods Table 1** for centre-specific reagent details.

### Electroporation mix preparation

Electroporation mixes were prepared essentially as previously reported^11–13,36^. Electroporation mixes consisted of Cas9 protein and sgRNA in RNP electroporation buffer (**Supplementary Methods Table 1**) at 2X the concentrations shown in **Supplementary Table S8**, incubated at 37°C or room temperature for 5-15 minutes, and placed on ice until electroporation. Immediately before electroporation, RNP was diluted with an equal volume of Opti-MEM (ThermoFisher 31985062). See **Supplementary Methods Table 1** for centre-specific reagent details.

### Generation of embryos by mating

C57BL/6N female mice, 3-6 weeks old, were injected with 5 IU/mouse of pregnant mare serum, followed 46-48 hr later with 5 IU/mouse of human chorionic gonadotropin. The females were then mated overnight with C57BL/6N males. Fertilized oocytes were collected from females with copulatory plugs the following morning at 0.5 days post-coitum (dpc). Oviducts were dissected and cumulus masses from these were released and treated with hyaluronidase. Fertilized 1-cell embryos were selected and maintained at 37°C in media prior to microinjection or electroporation.

### Microinjection of Cas9 reagents

The number of embryos injected and the injection route (pronuclear or cytoplasmic) for each experiment is in **Supplementary Table S8**. Pronuclear microinjections were performed following standard protocols^8,42^. Cytoplasmic injections were performed essentially as in ^9^. Visible movement of the cytoplasm indicated successful injection. Injected zygotes were transferred into pseudopregnant females (see **Supplementary Methods Table 1**) on the afternoon of the injection or after overnight culture (recorded for each production attempt in **Supplementary Table S8**), with 12-15 or 20-28 zygotes per unilateral or bilateral transfer into pseudopregnant females, respectively.

### Electroporation of Cas9 reagents

Electroporation was performed essentially as described^11–13,36^. At some centres, zygotes were briefly treated with Acid Tyrode’s solution (Sigma-Aldrich T1788). After acid treatment, embryos were rinsed at least 3 times with the final rinse in Opti-MEM. For electroporation, embryos were transferred into a 1:1 mixture of Cas9 RNP and Opti-MEM or Opti-MEM when RNP were formed in Opti-MEM. For each production attempt, electroporation pulses are in **Supplementary Table S8**. After electroporation the embryos were rinsed and transferred into pseudopregnant recipients the same day or after overnight culture (as recorded for each production attempt in **Supplementary Table S8**). Centre-specific details for buffers used are in **Supplementary Methods Table 1**.

### Genotyping

Genomic DNA was prepared from ear punch or tail biopsies of two- to three-week-old pups (see **Supplementary Methods Table 1** for reagents) using commercial kits or previously described protocols^43,44^. DNA was amplified by standard end-point PCR or quantitative PCR (qPCR). End-point PCR assays were designed to produce differently sized amplicons. To detect wild-type alleles, one primer was designed outside of the deletion target sequence and the second primer designed within the deletion target sequence such that amplicons are only produced from wild-type alleles. To detect deletion alleles, primers were designed to flank the predicted deletion junction. Amplification can result in two amplicons – a shorter amplicon representing the deletion allele and a larger amplicon representing the wild-type allele, if PCR conditions allow the amplification of the larger amplicon. Three-primer designs use a common primer outside of the deletion for both amplicons. PCR products were visualized using the Caliper LabChip GX system, QIAxcel Advanced, or agarose gel electrophoresis. Sequences are available upon request from the relevant production centre. In some cases, gene-specific ‘loss of WT allele’ (LoA) qPCR assays were designed to the region of the genome predicted to be deleted^45,46^. Deletion alleles will not amplify a product at the target site such that homozygous or hemizygous X-linked male deletions would have a copy number of 0, heterozygous a copy number of 1 and mosaic animals a copy number between 1 and 2 for autosomal alleles or between 0 and 1 in hemizygous X-linked alleles in males. These assays allowed estimation of the level of mosacism in founder animals. Mice showing the greatest loss of allele were selected for breeding to confirm germline transmission. Sequences for loss-of-allele assays are available upon request from the relevant production centres.

Once germline transmission was confirmed, mice were genotyped with either end-point PCR or probe-based LoA assays. Seep **Supplementary Methods Table 1** for centre-specific genotyping methods.

### Germline Transmission Test Breeding

Founders born from microinjection or electroporation experiments that carried the desired allele based on genotyping results were pair-mated to C57BL/6N mice. N1 pups were screened with the same genotyping assay used to identify founders. Deletion amplicons from deletion-positive N1 mice were subjected to Sanger sequencing (with or without subcloning) and other occasionally other quality control measures. **Copy number assessment:** When warranted, to assess whether the excised genomic fragment from deletion alleles re-inserted into the genome, DNA from N1 mice was purified using the NucleoSpin Tissue Kit (Machery-Nagel 740453) and subjected to digital droplet PCR (ddPCR) at The Centre for Applied Genomics (Toronto, Canada), the Mary Lyon Centre (Didcot, UK), or the Mouse Biology Program (University of California, Davis). The ddPCR assays were designed such that the amplification primers and probes were entirely contained within the target deletion fragment. For heterozygous N1 mice, a copy number equal to 1 (+/-0.2) was considered a pass; for hemizygous X-linked male mice, a copy number of 0 to 0.2 was considered a pass.

### Data download and filtering

A complete data set of Cas9-mediated mouse production attempts was downloaded on October 11, 2020 from the International Mouse Phenotyping Consortium production tracking database (formerly iMITS and now GenTaR; ‘Cas9 Micro-Injection Excel download’). This data included all Cas9-based production attempts as of that date. A production attempt was defined as the treatment of embryos to introduce Cas9 and guide RNAs to direct genome editing, subsequent embryo transfer, birth and screening of pups born from the embryo transfer, and subsequent breeding of mutant founders to obtain germline transmission of the desired edited allele. The data was filtered to remove attempts labeled as “private”, as “experimental”, or producing an allele other than a null allele, those with a status “Microinjection in Progress”, embryo transfer day of “Next Day”, none or >1000 embryos injected, incomplete information (e.g. number of founders not set, incomplete quality control information), and/or attempts that targeted non-protein coding genes. These data were further limited to attempts from production centres that had reported germline transmission for at least 50 unique genes for each of one or more of the analyzed methods (Cas9 mRNA pronuclear microinjection, Cas9 mRNA cytoplasmic injection, Cas9 RNP electroporation). This comprised the complete data set for analysis (**Supplementary Table 8**).

To define the set of unique gene experiments (i.e., each gene represented only once in the data set), the earliest attempt with germline transmission of the desired allele (Status = Genotype confirmed) for successful genes or the latest unsuccessful attempt (Status = Micro-injection aborted) was kept so that each gene was represented by a single attempt. If all attempts had a status of “Founder obtained”, the most recent was kept. However, if no attempts for a given gene were successful in this filtered dataset, the larger IMPC dataset was queried to see if a successful attempt existed in the pre-filtered dataset (e.g., at another IMPC production centre or as the result of technology development activities at a given centre). Successful production at another centre or through technology development activities could have resulted in aborting the production attempt in our filtered dataset, rather than failure of a complete experiment, or that technical issues rather than the parameters studied here resulted in failure, so these attempts were excluded from analysis.

For repeat attempt analysis, all attempts at the same production centre for genes that had more than one attempt were identified. This data set was then filtered to remove attempts in progress (Status = “Microinjection in progress” or “Founders obtained”). The remaining attempts were sorted in chronological order by microinjection [electroporation] date. If the first attempt for a given gene was successful, the set of attempts for that gene was removed from the repeat attempt analysis. Similarly, if an attempt was aborted within 6 weeks of a successful germline transmission attempt, it was removed since it may have been aborted because germline transmission had already been obtained, rather than having “failed”. Finally, if there was no GLT in any attempt at one centre, but successful GLT at another centre, the set of failed attempts was removed from the repeat dataset. The resulting data set comprised the repeat dataset (**Supplementary Table 3**).

### Data Annotation

Genes targeted for mouse line production attempts in **Supplementary Table 2** were annotated with derived parameters including bins for Cas9 mRNA and protein concentration, gRNA cut sites and predicted deletion sizes, percentage of embryos that survived to transfer of those treated (injected or electroporated), birth rate (number of pups born divided by embryos transferred), founder rate (number of founders born divided by embryos transferred), number of founders selected for breeding. Repeat attempts (**Supplementary Table 3**) were annotated with whether the Cas9 type (mRNA vs. protein), amount of Cas9, delivery of reagents (injection versus electroporation), or gRNA locations (sequences) changed between sequential attempts. All filtering and annotation of the data was performed in Python3.8.5 using packages numPy1.2.1^47^ and pandas1.2.2^48^.

Genes for each attempt were annotated (**Supplementary Table 4**) with their viability (as annotated at the IMPC – viable or homozygous lethal), human orthologs and cell essentiality of human orthologous genes^19^, embryonic expression (GEO GSE11224)^21^, length, GC content, number of CpG sites, and CpG percentage (**Supplementary Table 2**). The human orthologs’ probability of being loss-of-function intolerant (pLI) and observed / expected (oe) mutation rate was retrieved from gnomAD^22^. Additional annotations were added for analysis of biological variables affecting success. Annotation details are in **Supplementary Table 2**.

### Statistical Analysis

The primary outcomes were the founder rate and the germline transmission status. The founder rate had a right-skewed distribution with a range [0,0.5]. Hence, comparisons of the founder rate across different categories of biological or experimental factors were conducted using nonparametric tests. For pairwise comparisons, the Wilcoxon rank sum test^49^ was used and when there were more than two categories the Kruskal-Wallis test^50^ was employed. The biological factors considered in the comparisons were the gene essentiality (essential versus non-essential) and gene lethality (lethal versus non-lethal). The experimental factors were the delivery method (three categories), number of gRNAs used (2 versus 4), deletion size (six categories), and number of founders selected for breeding (four categories). Since the GLT status is binary (yes versus no), comparisons of the GLT rate (proportion of genes with GLT) across different categories of biological or experimental factors were performed using the Pearson chi-square test^51^. In the case of multiple pairwise comparisons, correction for multiple testing was done using Holm’s method^52^. Evaluation of success of repeated attempts was based on descriptive summaries, mainly calculation of relevant proportions. The assessment of the impact of changing experimental factors to the success of gene editing in repeated attempts was conducted using logistic regression with the GLT status as the binary response and changes in the delivery method (change versus no change), number of gRNAs used (decrease, no change, increase), deletion size (change versus no change) and number of founders selected for breeding (change versus no change) as categorical covariates. All statistical analyses were performed using the R statistical programing software^53^, along with the packages `ggplot2`^54^ for figures, `tidyverse`^55^ for data manipulations and `effects`^56,57^ for effect plots.

The general linear models of biological variables were fit using the glm function in the R 3.6.2 native stats package (https://rdocumentation.org/packages/stats/versions/3.6.2) using the factors in **Supplementary Table 4** and with founder rate as the dependent variable. All code can be found at https://github.com/The-Centre-for-Phenogenomics/IMPC-Cas9-Production.

## Author contributions

K.A.P., B.W., D.L., L.T., P.K., M-C. B., G.C., G.D., L.G., L.L., M.M., M.R., J.D.H., and L.M.J.N. designed alleles; D.L., E.R., P.K., A.C., G.C., B.D., G.D., M.G., L.G., L.L., K.L., I.L., M.M., S.A.M., and L.M.J.N. produced mouse lines or data; H.E. and P.M. developed software, databases and/or reporting tools; H.E., K.A.P., E.A., S.A.M., and L.M.J.N. analyzed data for the figures in the manuscript; B.W., D.L., L.T., P.K., M-C.B., D.A., A.B., R.B., S.B., A.C., M.D., F.D., L.G., K.L., I.L., M.M., A-M. M., C.M., H.P., R.R.-S., R.S., W.S., D.M., S.W., J.K.W., J.A.W., S.A.M., J.D.H., and L.M.J.N. co-supervised research; H.E., E.A., and L.M.J.N. wrote the manuscript. All authors reviewed and had the opportunity to comment on and edit the manuscript before submission.

## Acknowledgements

We thank all technical personnel at the IMPC production centres for their contributions. H.E., E.A., M.G., L.L., C.M., and L.M.J.N. were supported by Ontario Genomics and Genome Canada OGI-051, OGI-090, OGI-137 and the Canada Foundation for Innovation. M-C.B. and Y.H. were supported by the Université de Strasbourg, the CNRS, the INSERM and the ‘Investissements d’avenir’ programs (ANR-10-IDEX-0002-02, ANR-10-LABX-0030-INRT and ANR-10-INBS-07 PHENOMIN). A.C., G.C., M.M., L.T., and S.W. were supported by the Medical Research Council MC_UP_1502/3 International Mouse Phenotyping Consortium - building a functional catalogue of a mammalian genome. B.D., G.D., E.R., H.W.-J., D.A., A.B., R.R.-S., and W.S. were supported by the Wellcome Trust. P.M. and H.P. were supported by European Molecular Biology Laboratory core funding. P.K. and R.S. used services of the Czech Centre for Phenogenomics supported by the Czech Academy of Sciences RVO 68378050 and project LM2018126 Czech Centre for Phenogenomics provided by Ministry of Education, Youth and Sports of the Czech Republic, LM2015040 Czech Centre for Phenogenomics by MEYS, CZ.1.05/2.1.00/19.0395 Higher quality and capacity for transgenic models by MEYS and ERDF, CZ.1.05/1.1.00/02.0109 Biotechnology and Biomedicine Centre of the Academy of Sciences and Charles University in Vestec (BIOCEV) by MEYS and ERDF, CZ.02.1.01/0.0/0.0/16_013/0001789 Upgrade of the Czech Centre for Phenogenomics by MEYS and ESIF.

Research reported in this publication was supported by the NIH Common Fund, the Office of The Director and the National Human Genomic Research Institute of the National Institutes of Health (U42OD011174 and UMIHG006348 supported A.C., G.C., F.D., I.L., M.M., J.S., L.T., S.W., M.D., and J.D.H.; U42OD011175 and UM1OD023221 supported M.G., L.L., C.M., L.M.J.N., B.W., J.A.W., M.R., and K.L.; U42OD011185 and UM1OD023222 supported L.G., K.P., R.B., J.K.W., and S.A.M.; UM1HG006370 supported P.M., A.-M.M., and H.P.). The content is solely the responsibility of the authors and does not necessarily represent the official views of the National Institutes of Health.

## International Mouse Phenotyping Consortium (IMPC)

Shaheen Akhtar^8^, Alasdair J. Allan^9^, Susan Allen^9^, Philippe André^11^, Daniel Archer^9^, Sarah Atkins^9^, Ruth Avery^9^, Abdel Ayadi^11^, Daniel Barrett^8^, Tanya Beyetinova^8^, Toni Bell^9^, Melissa Berry^3^, Katharina Boroviak^8^, Joanna Bottomley^8^, Tim Brendler-Spaeth^8^, Ellen Brown^8^, Jonathan Burvill^8^, James Bussell^8^, Charis Cardeno^4^, Rebecca V. Carter^9^, Patricia Castellanos-Penton^1,2^, Skevoulla Christou^9^, Greg Clark^1,2^, Shannon Clarke^1,2^, James Cleak^9^, Ronnie Crawford^8^, Amie Creighton^1,2^, Maribelle Cruz^1^, Ozge Danisment^1,2^, Charlotte Davis^9^, Joanne Doran^8^, Valérie Erbs^11^, Qing Fan-Lan^1,2^, Rachel Fell^9^, He Feng^5^, Jean-Victor Fougerolle^11^, Alex Fower^9^, Gemma Frake^9^, Martin D. Fray^9^, Antonella Galli^8^, David Gannon^8^, Wendy J. Gardiner^9^, Angelina Gaspero^5^, Diane Gleeson^8^, Chris Godbehere^9^, Evelyn Grau^8^, Mark Griffiths^8^, Nicola Griggs^8^, Kristin Grimsrud^4^, Sarah Hazeltine^4^, Marie Hutchison^9^, Catherine Ingle^8^, Vivek Iyer^8^, Kayla Jager^4^, Joanna Joeng^1,2^, Susan Kales^3^, Perminder Kaur^4^, Janet Kenyon^9^, Jana Kopkanova^10^, Christelle Kujath^11^, Helen Kundi^8^, Peter Kutny^3^, Valerie Laurin^1,2^, Sandrine Lejeay^11^, Christopher Lelliott^8^, Jorik Loeffler^9^, Romain Lorentz^11^, Christopher V. McCabe^9^, Elke Malzer^9^, Peter Matthews^16^, Ryea Maswood^8^, Matthew McKay^3^, Terrence Meehan^16^, David Melvin^8^, Alison Murphy^8^, Asif Nakhuda^8^, Amit Patel^1^, Ilya Paulavets^8^, Guillaume Pavlovic^11^, Ashley Pawelka^5^, Fran J. Pike^9^, Radka Platte^8^, Peter D. Price^9^, Kiran Rajaya^5^, Shalini Reddy^8^, Whitney Rich^5^, Barry Rosen^8^, Victoria Ross^8^, Mark Ruhe^4^, Luis Santos^15^, Laurence Schaeffer^11^, Alix Schwiening^8^, Mohammed Selloum^11^, Debarati Sethi^8^, Jan R. Sidiangco^9^, Caroline Sinclair^8^, Elodie Sins^8^, Gillian Sleep^1,2^, Tania Sorg^11^, Becky Starbuck^9^, Michelle Stewart^9^, Holly Swash^9^, Mark Thomas^8^, Sandra Tondat^1^, Rachel Urban^3^, Jana Urbanova^8^, Susan Varley^9^, Dominque Von Schiller^8^, Hannah Wardle-Jones^8^, Lauren Weavers^8^, Michael Woods^8^

## Notes

https://github.com/The-Centre-for-Phenogenomics/IMPC-Cas9-Production

https://www.i-dcc.org/imits/

### Competing Interest Statement

The authors have declared no competing interest.

### Summary of Updates

The core analyses presented in this manuscript remain the same, but we have provided more explicit recommendations about optimizing genome editing experiments in mice. In addition, the information about the larger resource has been removed and will be presented in a separate manuscript in the future.

## References

1 Lloyd, K. C. K. et al. The Deep Genome Project. Genome Biol 21, 18, doi:10.1186/s13059-020-1931-9 (2020).

2 Birling, M. C. et al. A resource of targeted mutant mouse lines for 5,061 genes. Nat Genet 53, 416–419, doi:10.1038/s41588-021-00825-y (2021).

3 Bradley, A. et al. The mammalian gene function resource: the international knockout mouse consortium. Mamm Genome 23, 580–586, doi:10.1007/s00335-012-9422-2 (2012).

4 Popp, M. W. & Maquat, L. E. The dharma of nonsense-mediated mRNA decay in mammalian cells. Mol Cells 37, 1–8, doi:10.14348/molcells.2014.2193 (2014).

5 Smits, A. H. et al. Biological plasticity rescues target activity in CRISPR knock outs. Nat Methods 16, 1087–1093, doi:10.1038/s41592-019-0614-5 (2019).

6 Lalonde, S. et al. Frameshift indels introduced by genome editing can lead to in-frame exon skipping. PLoS One 12, e0178700, doi:10.1371/journal.pone.0178700 (2017).

7 Mou, H. et al. CRISPR/Cas9-mediated genome editing induces exon skipping by alternative splicing or exon deletion. Genome Biol 18, 108, doi:10.1186/s13059-017-1237-8 (2017).

8 Behringer, R. R., Gertsenstein, M., Nagy, K. & Nagy, A. Manipulating the Mouse Embryo: A Laboratory Manual. 4th edn, (Cold Spring Harbor Laboratory Press, 2014).

9 Doe, B., Brown, E. & Boroviak, K. Generating CRISPR/Cas9-Derived Mutant Mice by Zygote Cytoplasmic Injection Using an Automatic Microinjector. Methods Protoc 1, doi:10.3390/mps1010005 (2018).

10 Gertsenstein, M. & Nutter, L. M. J. Production of knockout mouse lines with Cas9. Methods 191, 32–43, doi:10.1016/j.ymeth.2021.01.005 (2021).

11 Wang, W. et al. Delivery of Cas9 Protein into Mouse Zygotes through a Series of Electroporation Dramatically Increases the Efficiency of Model Creation. Journal of genetics and genomics = Yi chuan xue bao 43, 319–327, doi:10.1016/j.jgg.2016.02.004 (2016).

12 Kaneko, T., Sakuma, T., Yamamoto, T. & Mashimo, T. Simple knockout by electroporation of engineered endonucleases into intact rat embryos. Scientific reports 4, 6382, doi:10.1038/srep06382 (2014).

13 Modzelewski, A. J. et al. Efficient mouse genome engineering by CRISPR-EZ technology. Nat Protoc 13, 1253–1274, doi:10.1038/nprot.2018.012 (2018).

14 Peterson, K. A. et al. Whole genome analysis for 163 gRNAs in Cas9-edited mice reveals minimal off-target activity. Commun Biol 6, 626, doi:10.1038/s42003-023-04974-0 (2023).

15 Iyer, V. et al. Off-target mutations are rare in Cas9-modified mice. Nat Methods 12, 479, doi:10.1038/nmeth.3408 (2015).

16 Anderson, K. R. et al. CRISPR off-target analysis in genetically engineered rats and mice. Nat Methods 15, 512–514, doi:10.1038/s41592-018-0011-5 (2018).

17 Willi, M., Smith, H. E., Wang, C., Liu, C. & Hennighausen, L. Mutation frequency is not increased in CRISPR-Cas9-edited mice. Nat Methods 15, 756–758, doi:10.1038/s41592-018-0148-2 (2018).

18 Lanza, D. G. et al. Comparative analysis of single-stranded DNA donors to generate conditional null mouse alleles. BMC biology 16, 69, doi:10.1186/s12915-018-0529-0 (2018).

19 Cacheiro, P. et al. Human and mouse essentiality screens as a resource for disease gene discovery. Nature communications 11, 655, doi:10.1038/s41467-020-14284-2 (2020).

20 Ring, N. et al. A mouse informatics platform for phenotypic and translational discovery. Mamm Genome 26, 413–421, doi:10.1007/s00335-015-9599-2 (2015).

21 Barrett, T., et al. NCBI GEO: archive for functional genomics data sets--update. Nucleic Acids Res 41, D991–995, doi:10.1093/nar/gks1193 (2013).

22 Karczewski, K. J. et al. The mutational constraint spectrum quantified from variation in 141,456 humans. Nature 581, 434–443, doi:10.1038/s41586-020-2308-7 (2020).

23 (McKusick-Nathans Institute of Genetic Medicine, Johns Hopkins University (Baltimore, MD), 2020).

24 Scavizzi, F. et al. Blastocyst genotyping for quality control of mouse mutant archives: an ethical and economical approach. Transgenic Res 24, 921–927, doi:10.1007/s11248-015-9897-1 (2015).

25 Economides, A. N. et al. Conditionals by inversion provide a universal method for the generation of conditional alleles. Proc Natl Acad Sci U S A 110, E3179–3188, doi:10.1073/pnas.1217812110 (2013).

26 Nagy, A. Cre recombinase: the universal reagent for genome tailoring. Genesis 26, 99–109, doi:10.1002/(SICI)1526-968X(200002)26:2<99::AID-GENE1>3.0.CO;2-B [pii] (2000).

27 Cacheiro, P. et al. Mendelian gene identification through mouse embryo viability screening. Genome Med 14, 119, doi:10.1186/s13073-022-01118-7 (2022).

28 Russell, W. M. S. & Burch, R. L. The Principles of Humane Experimental Technique. (Methuen, 1959).

29 Popp, M. W. & Maquat, L. E. Organizing principles of mammalian nonsense-mediated mRNA decay. Annu Rev Genet 47, 139–165, doi:10.1146/annurev-genet-111212-133424 (2013).

30 Ran, F. A. et al. Genome engineering using the CRISPR-Cas9 system. Nat Protoc 8, 2281–2308, doi:10.1038/nprot.2013.143 (2013).

31 Hodgkins, A. et al. WGE: a CRISPR database for genome engineering. Bioinformatics 31, 3078–3080, doi:10.1093/bioinformatics/btv308 (2015).

32 Haeussler, M. et al. Evaluation of off-target and on-target scoring algorithms and integration into the guide RNA selection tool CRISPOR. Genome Biol 17, 148, doi:10.1186/s13059-016-1012-2 (2016).

33 Peterson, K. A. et al. CRISPRtools: a flexible computational platform for performing CRISPR/Cas9 experiments in the mouse. Mamm Genome 28, 283–290, doi:10.1007/s00335-017-9681-z (2017).

34 Labun, K., Montague, T. G., Gagnon, J. A., Thyme, S. B. & Valen, E. CHOPCHOP v2: a web tool for the next generation of CRISPR genome engineering. Nucleic Acids Res 44, W272–276, doi:10.1093/nar/gkw398 (2016).

35 Elrick, H., et al. FORCAST: a fully integrated and open source pipeline to design Cas-mediated mutagenesis experiments. bioRxiv, 2020.2004.2021.053090, doi:10.1101/2020.04.21.053090 (2020).

36 Gertsenstein, M. & Nutter, L. M. J. Engineering point mutant and epitope-tagged alleles in mice using Cas9 RNA-guided nuclease. Curr Protoc Mouse Biol 8, 28–53, doi:10.1002/cpmo.40 (2018).

37 Bassett, A. R., Tibbit, C., Ponting, C. P. & Liu, J. L. Highly efficient targeted mutagenesis of Drosophila with the CRISPR/Cas9 system. Cell reports 4, 220–228, doi:10.1016/j.celrep.2013.06.020 (2013).

38 Cong, L. et al. Multiplex genome engineering using CRISPR/Cas systems. Science 339, 819–823, doi:10.1126/science.1231143 (2013).

39 Mali, P. et al. RNA-guided human genome engineering via Cas9. Science 339, 823–826, doi:10.1126/science.1232033 (2013).

40 Hwang, W. Y. et al. Efficient genome editing in zebrafish using a CRISPR-Cas system. Nat Biotechnol 31, 227–229, doi:10.1038/nbt.2501 (2013).

41 Mianne, J. et al. Correction of the auditory phenotype in C57BL/6N mice via CRISPR/Cas9-mediated homology directed repair. Genome Med 8, 16, doi:10.1186/s13073-016-0273-4 (2016).

42 Gardiner, W. J. & Teboul, L. Overexpression transgenesis in mouse: pronuclear injection. Methods Mol Biol 561, 111–126, doi:10.1007/978-1-60327-019-9_8 (2009).

43 Truett, G. E. et al. Preparation of PCR-quality mouse genomic DNA with hot sodium hydroxide and tris (HotSHOT). Biotechniques 29, 52, 54, doi:10.2144/00291bm09 (2000).

44 Green, M. R. & Sambrook, J. Molecular Cloning: A Laboratory Manual. Vol. 3 (Cold Spring Harbor Laboratory Press, 2012).

45 Ryder, E. et al. Molecular characterization of mutant mouse strains generated from the EUCOMM/KOMP-CSD ES cell resource. Mamm Genome 24, 286–294, doi:10.1007/s00335-013-9467-x (2013).

46 Mianne, J. et al. Analysing the outcome of CRISPR-aided genome editing in embryos: Screening, genotyping and quality control. Methods 121-122, 68–76, doi:10.1016/j.ymeth.2017.03.016 (2017).

47 Harris, C. R. et al. Array programming with NumPy. Nature 585, 357–362, doi:10.1038/s41586-020-2649-2 (2020).

48 McKinney, W. in 9th Python in Science Conference. (eds S. van der Walt & J. Millman) 50–61.

49 Wilcoxon, F. Individual comparisons by ranking methods. Biometrics Bulletin 1, 80–83, doi:10.2307/3001968 (1945).

50 Kruskal, W. H. & Wallis, W. A. Use of Ranks in One-Criterion Variance Analysis. Journal of the American Statistical Association 47, 583–621, doi:10.1080/01621459.1952.10483441 (1952).

51 Pearson, K. X. On the criterion that a given system of deviations from the probable in the case of a correlated system of variables is such that it can be reasonably supposed to have arisen from random sampling. The London, Edinburgh, and Dublin Philosophical Magazine and Journal of Science 50, 157–175, doi:10.1080/14786440009463897 (1900).

52 Holm, S. A Simple Sequentially Rejective Multiple Test Procedure. Scandinavian Journal of Statistics 6, 65–70, doi:10.2307/4615733 (1979).

53 Team, R. C. in R Foundation for Statistical Computing, (2021).

54 Wickham, H. ggplot2: Elegant Graphics for Data Analysis. (Springer-Verlag, 2016).

55 Wickham, H. et al. Welcome to the Tidyverse. The Journal of Open Source Software 4, 1686, doi:10.21105/joss.01686 (2019).

56 Fox, J. Effect Displays in R for Generalised Linear Models. 2003 8, 27, doi:10.18637/jss.v008.i15 (2003).

57 Fox, J. & Weisberg, S. An R companion to applied regression. Third edition edn, (SAGE, 2019).

