## Supplemental Material for "Impact of Essential Genes on the Success of Genome Editing Experiments Generating 3,313 New Genetically Engineered Mouse Lines"

Elrick *et al*.

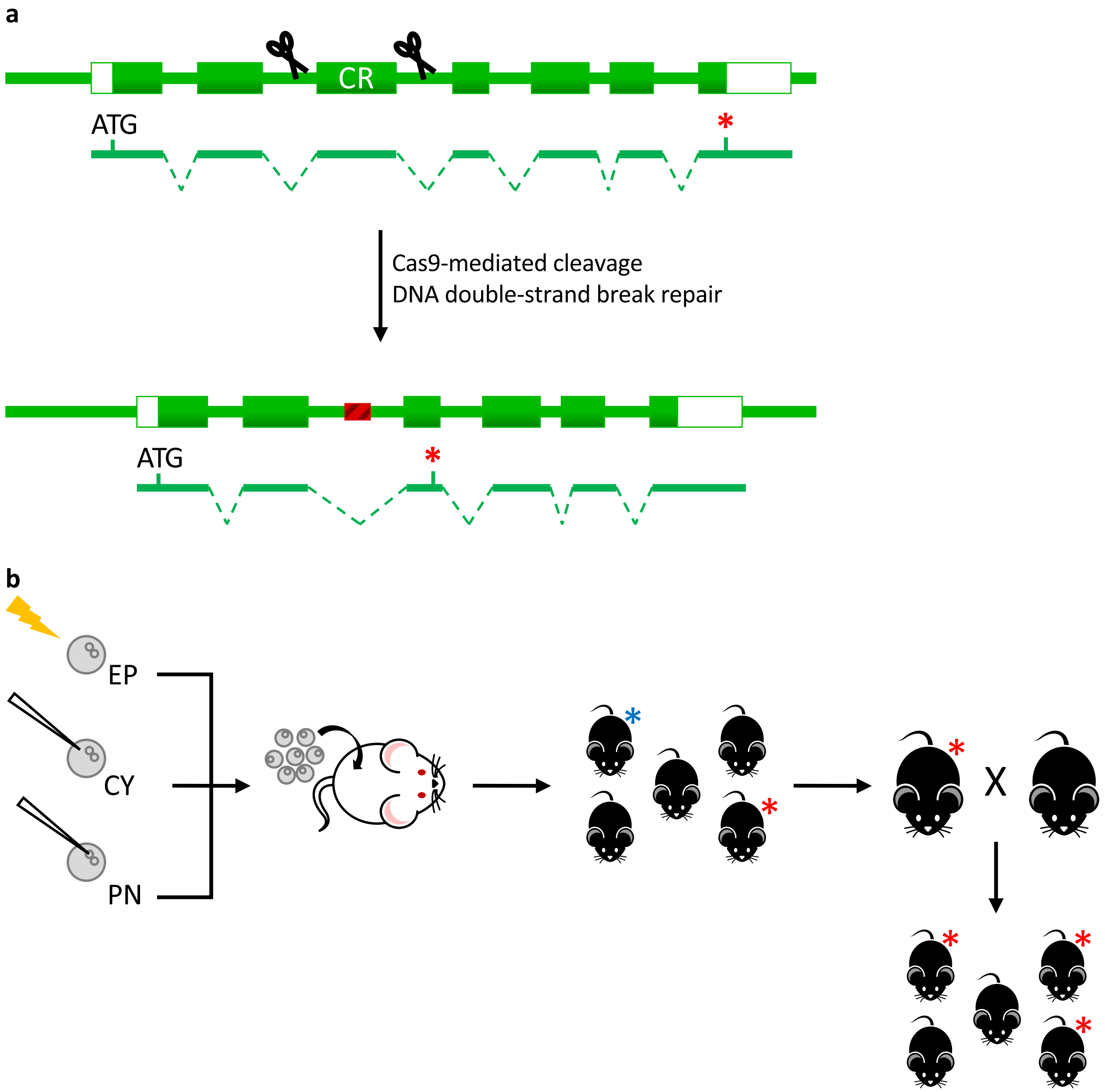


**Supplementary Figure 1**. Schematic of null allele mouse line production. **a**. Allele Design. A critical region (CR) was identified that contains one or more exons present in all annotated fulllength protein-coding transcripts and when deleted would introduce a frameshift and premature stop codon in the first half of the open reading frame but at least 33 amino acids downstream of the translation start codon. In most cases, these transcripts are predicted to be targeted for nonsense-mediated decay. Intron sequences flanking the CR were examined for specific Cas9 protospacer sequences. Cas9 guide RNA specificity was gauged by the absence of predicted off-target sites with fewer than three mismatches and/or a specificity score >65. At some centres, specificity parameters included that off-target sites must have at least one mismatch in the seed region (11-bp immediately 5' to the protospacer adjacent motif) of the guide RNA. **b**. Mouse Production. Cas9 reagents were introduced into groups of zygotes by electroporation (EP) or pronuclear (PN) or cytoplasmic (CY) microinjection. Treated zygotes were transferred to pseudopregnant recipient for gestation and birth. Born pups were genotyped for the presence of the deletion allele, either by real-time PCR or droplet digital PCR to quantify the copies of the deleted region or by end-point PCR to detect an amplicon consistent with the deletion of the critical region. Founders were bred to wild-type C57BL/6N mice and N1 pups were screened for the presence of the deletion. Sanger sequencing of a PCR amplicon spanning the deletion confirmed the sequence of the deletion junction and at least 100-bp of DNA flanking either side of the deletion. N1 pups with the same allele sequence were used to establish the putative null mutant mouse line.

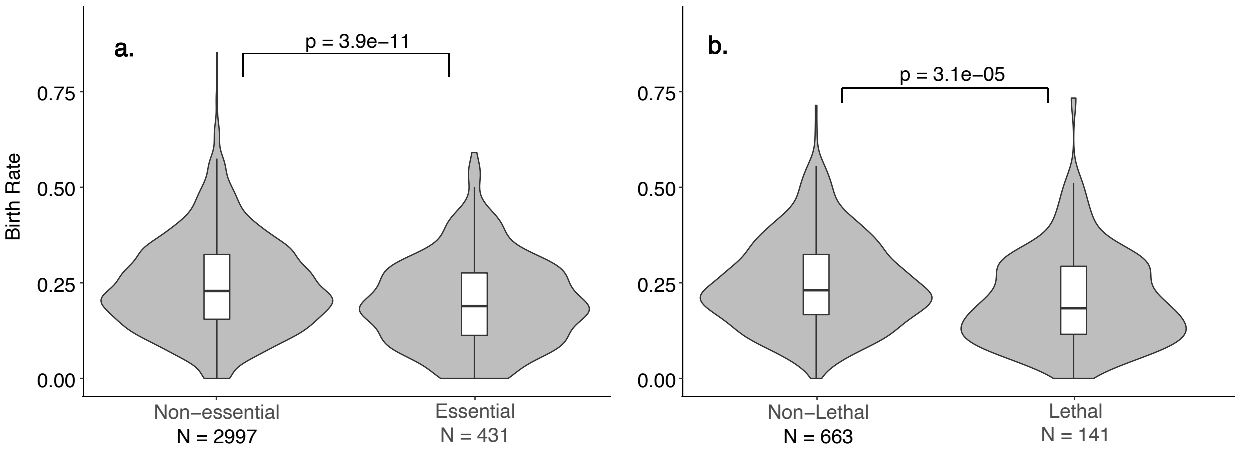


**Supplementary Figure 2**. Birth rates from experiments generating null alleles in (a) non-essential and essential genes as well as (b) non-lethal and lethal genes.

**Supplementary Table 1.** Experiments included in meta-analysis by production centre

| | BCM | CCP | HAR | ICS | JAX | TCP | UCD | WTSI | Total |
|---|---|---|---|---|---|---|---|---|---|
| Number of experiments | 679 | 339 | 413 | 60 | 1430 | 386 | 701 | 468 | 4473 |
| Number of genes | 585 | 278 | 305 | 60 | 1340 | 341 | 683 | 383 | 3973 |
| %Repeated experiments | 25.8 | 32.7 | 41.0 | 0.0 | 12.4 | 22.5 | 5.1 | 31.2 | 20.1 |
| %Repeated genes | 13.8 | 18.0 | 20.7 | 0.0 | 6.6 | 12.3 | 2.6 | 15.9 | 10.1 |

BCM, Baylor College of Medicine; CCP, Czech Centre for Phenogenomics; HAR, MRC Harwell; ICS, Institut de Clinque des Souris; JAX, The Jackson Laboratory; TCP, The Centre for Phenogenomics; UCD, University of California, Davis, Mouse Biology Program; WTSI, Wellcome Trust Sanger Institute.

Supplementary Table 2 – ST2_UniqueGeneCas9Attempts.xlsx

Supplementary Table 3 – ST3_RepeatedGeneCas9Attempts.xlsx

Supplementary Table 4 – ST4_Cas9_GeneAnnotations.xlsx

**Supplementary Table 5**. General linear model (GLM) output
Observations: 3209
Dependent Variable: Founders
Type: Linear regression

**GLM without Essentiality**

| | Odds Ratio | Standard Error | t-value | p-value |
|---|---|---|---|---|
| (Intercept) | 2.52 | 0.0251 | 36.882 | $2.0x10^{-16}$ |
| Embryonic Expression | 0.95 | 0.0210 | -2.552 | 0.011* |
| pLI | 0.97 | 0.0193 | -1.638 | 0.101 |
| oe | 1.00 | 0.0242 | -0.016 | 0.987 |
| Chromosomal Location | 1.01 | 0.0197 | 0.458 | 0.647 |
| Acetylated | 0.98 | 0.0129 | -1.301 | 0.193 |
| Methylated | 1.03 | 0.0173 | 1.486 | 0.137 |
| Gram Positive Stain | 1.00 | 0.0112 | -0.078 | 0.938 |
| OMIM Annotation | 0.99 | 0.0160 | -0.821 | 0.412 |

**GLM with Essentiality**

| | Odds Ratio | Standard Error | t-value | p-value |
|---|---|---|---|---|
| (Intercept) | 2.52 | 0.025 | 37.146 | $2.22x10^{-16}$ |
| Essential | 0.89 | 0.017 | -6.726 | $2.06x10^{-11}$ |
| Embryonic Expression | 0.97 | 0.021 | -1.330 | 0.219 |
| pLI | 0.97 | 0.019 | -1.333 | 0.183 |
| oe | 1.00 | 0.024 | -1.142 | 0.887 |
| Chromosomal Location | 1.01 | 0.020 | 0.449 | 0.653 |
| Acetylated | 0.99 | 0.013 | -0.755 | 0.450 |
| Methylated | 1.01 | 0.017 | 0.872 | 0.383 |
| Gram Positive Stain | 1.00 | 0.011 | 0.136 | 0.891 |
| OMIM Annotation | 0.98 | 0.016 | -1.227 | 0.220 |

| MODEL FIT: Without Essentiality | MODEL FIT: With Essentiality |
|---|---|
| $\chi^2(8) = 2.54$, p = 0.00 | $\chi^2(9) = 6.85$, p = 0.00 |
| Pseudo-$R^2$ (Cragg-Uhler) = 0.02 | Pseudo-$R^2$ (Cragg-Uhler) = 0.06 |
| Pseudo-$R^2$ (McFadden) = 0.02 | Pseudo-$R^2$ (McFadden) = 0.04 |
| AIC = 1619.49, BIC = 1680.22 | AIC = 1576.43, BIC = 1643.24 |

**Supplementary Table 6.** The number of genes across chromosomes and their distribution based on essentiality

| Chromosome | No. Genes | Essential | Non-essential | Unknown | %known* |
|---|---|---|---|---|---|
| 1 | 251 | 24 | 205 | 22 | 10.5% |
| 2 | 277 | 30 | 240 | 7 | 11.1% |
| 3 | 212 | 21 | 173 | 18 | 10.8% |
| 4 | 212 | 23 | 180 | 9 | 11.3% |
| 5 | 254 | 38 | 200 | 16 | 16.0% |
| 6 | 192 | 15 | 155 | 22 | 8.8% |
| 7 | 311 | 36 | 237 | 38 | 13.2% |
| 8 | 208 | 28 | 173 | 7 | 13.9% |
| 9 | 251 | 26 | 206 | 19 | 11.2% |
| 10 | 170 | 24 | 132 | 14 | 15.4% |
| 11 | 262 | 31 | 216 | 15 | 12.6% |
| 12 | 129 | 17 | 99 | 13 | 14.7% |
| 13 | 145 | 19 | 106 | 20 | 15.2% |
| 14 | 144 | 12 | 125 | 7 | 8.8% |
| 15 | 150 | 19 | 120 | 11 | 13.7% |
| 16 | 126 | 15 | 106 | 5 | 12.4% |
| 17 | 172 | 24 | 131 | 17 | 15.5% |
| 18 | 112 | 15 | 85 | 12 | 15.0% |
| 19 | 131 | 14 | 108 | 9 | 11.5% |
| X | 159 | 0 | 0 | 159 | nd |
| Y | 1 | 0 | 0 | 1 | nd |
| **TOTAL** | **3869** | **431** | **2997** | **441** | **12.6%** |

*known = genes for which essentiality of human ortholog has been reported

**Supplementary Table 7.** Pairwise comparisons of the proportion of essential genes by experimental parameter

| Experimental Parameter | Comparison (left vs. right) | % left | % right | p-value* |
|---|---|---|---|---|
| Delivery method | Cytoplasmic injection vs. electroporation | 11.0% | 14.2% | 0.025 |
| | Cytoplasmic vs. pronuclear injection | 11.0% | 7.2% | 0.077 |
| | Electroporation vs. pronuclear injection | 14.2% | 7.2% | 0.004 |
| No. guides | 2 guides vs. 4 guides | 12.8% | 12.7% | 0.986 |
| Deletion size | [35,360] vs. [360,490] | 13.8% | 11.1% | 1.000 |
| | [35,360] vs. [490,640] | 13.8% | 12.6% | 1.000 |
| | [35,360] vs. [640,880] | 13.8% | 13.6% | 1.000 |
| | [35,360] vs. [880,1400] | 13.8% | 8.6% | 0.095 |
| | [35,360] vs. [1400+] | 13.8% | 15.9% | 1.000 |
| | [360,490] vs. [490,640] | 11.1% | 12.6% | 1.000 |
| | [360,490] vs. [640,880] | 11.1% | 13.6% | 1.000 |
| | [360,490] vs. [880,1400] | 11.1% | 8.6% | 1.000 |
| | [360,490] vs. [1400+] | 11.1% | 15.9% | 0.274 |
| | [490,640] vs. [640,880] | 12.6% | 13.6% | 1.000 |
| | [490,640] vs. [880,1400] | 12.6% | 8.6% | 0.395 |
| | [490,640] vs. [1400+] | 12.6% | 15.9% | 1.000 |
| | [640,880] vs. [880,1400] | 13.6% | 8.6% | 0.124 |
| | [640,880] vs. [1400+] | 13.6% | 15.9% | 1.000 |
| | [880,1400] vs. [1400+] | 8.6% | 15.9% | 0.004 |
| No. founders bred | 1 vs. 2 | 10.1% | 13.7% | 0.132 |
| | 1 vs. 3 | 10.1% | 9.4% | 1.000 |
| | 1 vs. 4 | 10.1% | 9.6% | 1.000 |
| | 2 vs. 3 | 13.7% | 9.4% | 0.132 |
| | 2 vs. 4+ | 13.7% | 9.6% | 0.132 |
| | 3 vs. 4+ | 9.4% | 9.6% | 1.000 |

*Chi square test for proportions

Supplementary Table 8 - ST8_Cas9_AllAttempts.xlsx
