## Supplemental Methods for "Impact of Essential Genes on the Success of Genome Editing Experiments Generating 3,313 New Genetically Engineered Mouse Lines"

Elrick et al.

**Supplemental Methods Table 2.** Main Dataset Annotation Table

| Annotation | File | Source |
|---|---|---|
| Protein Coding Genes | protein-coding-mgi_14-02-2021.csv | Mouse Genome Database (MGD) at the Mouse Genome Informatics website, The Jackson Laboratory, Bar Harbor, Maine. World Wide Web (URL: http://www.informatics.jax.org). |
| Viability Consensus | IMPC-viability_release-13.0.csv | IMPC[1] |
| Cellular Essential | HumanEssentiality_MouseOrthologs.xlsx | [2] |
| Human Ortholog | Human_mouse_orthologues_for_IMPC.xlsx | [3] |
| Ortholog Relationship (human-to-mouse_mouse-to-human) | Human_mouse_orthologues_for_IMPC.xlsx | [3] |
| Embryonic Expression | GPL1261-56135.txt, GSE11224_series_matrix.txt | [4] NCBI GEO database[5], accession GSE11224 |
| pLI of Orthologs | gnomad_v2_1_1_lof_metrics_by_gene.xlsx | [6] |
| oe of Orthologs | gnomad_v2_1_1_lof_metrics_by_gene.xlsx | [6] |

**Supplemental Methods Table 3.** Generalized Linear Model Analysis Annotation Table

| Annotation | File | Source |
|---|---|---|
| Acetylated and Methylated Regions | mm10_H3K27ac_annotatedpeaks.tsv | We downloaded the call sets from the ENCODE portal[7] (https://www.encodeproject.org/) with the following identifiers: ENCSR000CDE and ENCSR000CFN and annotated them with HOMER[8] |
| Human Ortholog OMIM Entry | omim_to_ensembl.tsv | [9]URL: https://omim.org/ |
| Cytogenic Banding (Gram Positive) | cytoBand.txt | [10] |
| Chromosome Sizes (for relative position calculations) | mm10.chrom.sizes | |

**Supplemental Methods Table 4.** Factors affecting the dependent variable founder rate in the generalized linear model.

| Annotation | Type | Source |
|---|---|---|
| Cellular Essential Gene | Categorical (two levels) | [3] |
| Embryonic Expression | Continuous | GEO dataset GSE11224 |
| pLI of Human Ortholog | Continuous | Gnomad version 2.1.1 |
| oe of Human Ortholog | Continuous | Gnomad version 2.1.1 |
| Relative Chromosomal Position | Continuous | Cut-site divided by chromosome length as defined in [ucsc link] |
| Methylated Region | Categorical (two levels) | GEO dataset ENCSR000CDE, annotated with homer[8] |
| Acetylated Region | Categorical (two levels) | GEO dataset ENCSR000CFN annotated with homer[8] |
| Gram Positive | Categorical (two levels) | Cytogenic band file for mm10 |
| Human Ortholog Present in OMIM | Categorical (two levels) | Downloaded from OMIM March 10th, 2020 |

### References


1 Dickinson, M. E. *et al.* High-throughput discovery of novel developmental phenotypes. *Nature* **537**, 508-514, doi:10.1038/nature19356 (2016).

2 Tsherniak, A. *et al.* Defining a Cancer Dependency Map. *Cell* **170**, 564-576 e516, doi:10.1016/j.cell.2017.06.010 (2017).

3 Cacheiro, P. *et al.* Human and mouse essentiality screens as a resource for disease gene discovery. *Nature communications* **11**, 655, doi:10.1038/s41467-020-14284-2 (2020).

4 Knox, K. & Baker, J. C. Genomic evolution of the placenta using co-option and duplication and divergence. *Genome Res* **18**, 695-705, doi:10.1101/gr.071407.107 (2008).

5 Barrett, T. *et al.* NCBI GEO: archive for functional genomics data sets--update. *Nucleic Acids Res* **41**, D991-995, doi:10.1093/nar/gks1193 (2013).

6 Karczewski, K. J. *et al.* The mutational constraint spectrum quantified from variation in 141,456 humans. *Nature* **581**, 434-443, doi:10.1038/s41586-020-2308-7 (2020).

7 Sloan, C. A. *et al.* ENCODE data at the ENCODE portal. *Nucleic Acids Res* **44**, D726-732, doi:10.1093/nar/gkv1160 (2016).

8 Heinz, S. *et al.* Simple combinations of lineage-determining transcription factors prime cis-regulatory elements required for macrophage and B cell identities. *Mol Cell* **38**, 576-589, doi:10.1016/j.molcel.2010.05.004 (2010).

9 (McKusick-Nathans Institute of Genetic Medicine, Johns Hopkins University (Baltimore, MD), 2020).

10 Navarro Gonzalez, J. *et al.* The UCSC Genome Browser database: 2021 update. *Nucleic Acids Res* **49**, D1046-D1057, doi:10.1093/nar/gkaa1070 (2021).
